# Testing the Spatiotemporal Predictions of Global Neuronal Workspace Theory and Integrated Information Theory

**DOI:** 10.64898/2026.07.16.738188

**Authors:** Kavindu H. Bandara, Elise G. Rowe, Marta I. Garrido

## Abstract

The neural mechanisms underlying conscious perception remain contested, with global neuronal workspace theory (GNWT) and integrated information theory (IIT) offering divergent predictions about the spatiotemporal dynamics of conscious processing. The Cogitate Consortium (2025) recently conducted a large-scale adversarial collaboration to arbitrate between GNWT and IIT; however, some of their analyses yielded mixed findings. As the original study relied on functional connectivity, it could not assess the directed neural influences required to directly test the mechanistic claims of these theories. Here, we re-examined the Cogitate Consortium’s MEG dataset (N = 100) using dynamic causal modelling (DCM) to test the effective connectivity profiles predicted by each theory during task-irrelevant face perception. We adopted a split discovery-validation procedure and an expanding time-window approach to track the temporal evolution of prototypical networks for each theory. Specifically, we evaluated GNWT’s prediction of phasic prefrontal “ignition” by testing the necessity of PFC feedback to category-selective areas at stimulus onset and offset, and then tested IIT’s prediction of sustained posterior integration by testing extrinsic feedback and intrinsic connectivity within content-specific regions. We found support for both GNWT predictions of prefrontal ignition at stimulus onset and offset; noting that the latter was not previously observed in the original Cogitate analyses. For IIT, whilst feedback connectivity between the fusiform gyrus and primary visual cortex was not sustained throughout the stimulus duration, intrinsic fusiform gyrus connectivity was broadly consistent with IIT’s prediction of sustained processing within a content-specific substrate. However, a post-hoc validation analysis comparing face- and object-specific DCM networks did not yield strong evidence for face selectivity at any time window (pp< 0.95), warranting a conservative interpretation of the IIT findings in particular.

---

Despite a surge of interest and accumulating empirical data in the science of consciousness, researchers have failed to conclusively arbitrate between the many theories of consciousness that now exist (Seth & Bayne, 2022). This is due, in part, to the fact that different theoretical models of consciousness have developed in isolation and rarely receive direct comparison (Yaron et al. 2022). To address this, the Cogitate Consortium et al. (2025) recently conducted a large-scale adversarial collaboration directly comparing two of the most prominent contemporary theories of consciousness: global neuronal workspace theory (GNWT; Dehaene & Naccache, 2001; Mashour et al., 2020) and integrated information theory (IIT; Tononi et al., 2016; Albantakis et al., 2023). However, since the original study only considered the data in terms of functional connectivity, it could not test the directed influence of different regions on each other. In this chapter, we address this gap by re-examining this dataset using dynamic causal modelling (DCM) to formally test putative transient prefrontal connectivity as predicted by GNWT versus sustained posterior integration predicted by IIT, during conscious perception.

To summarise the key theoretical predictions of GNWT, the theory postulates that consciousness is characterised by a late-stage non-linear response (termed an ignition) occurring within 300-500ms post-stimulus onset and offset occurring in a large scale frontoparietal network (see Figure 1). This is because an onset and offset are both considered an update in the percept and hence the offset too should elicit an ignition as it is essentially the “onset” of a blank screen. Further, the theory highlights the importance of feedback connections during the ignition phase, important for the amplification and broadcasting of conscious information (Dehaene et al., 2003; Dehaene & Changeux, 2005; Mashour et al., 2020).

**Figure 1.**
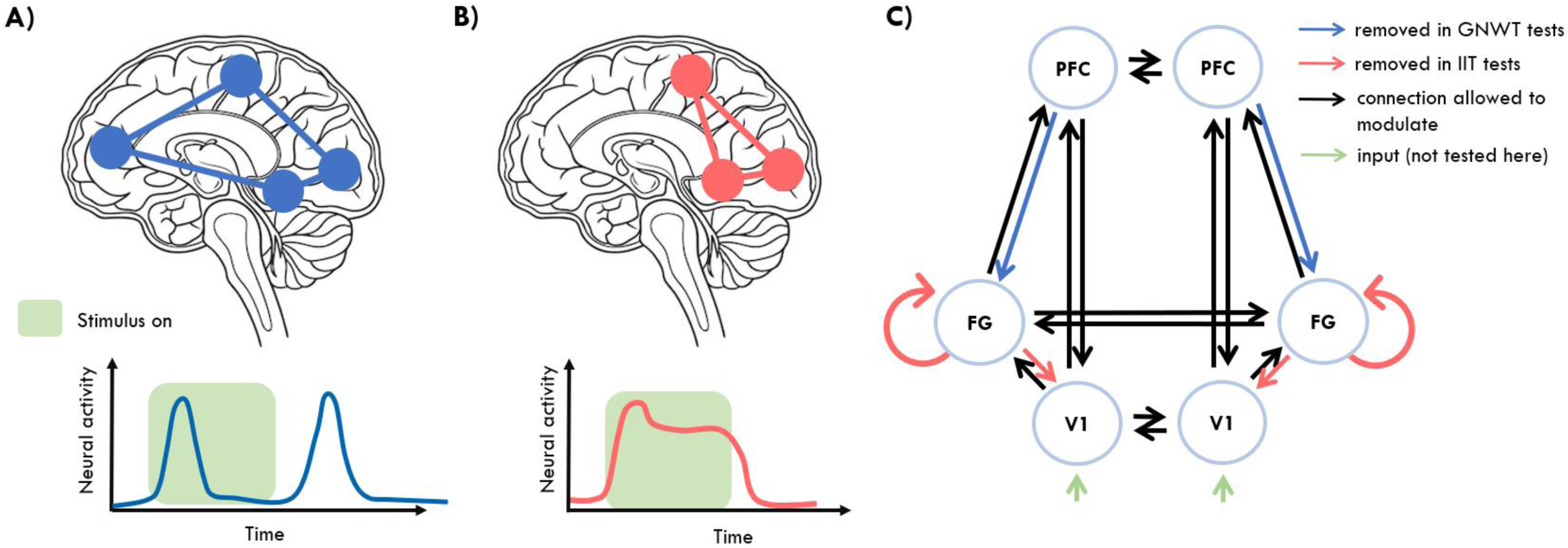
Divergent spatiotemporal predictions of GNWT and IIT and study design. **(A)** GNWT predicts consciousness is associated with a “global ignition” event in a frontoparietal network (the global workspace). Temporally, this manifests as a late, phasic burst of activity at both stimulus onset and offset (∼300ms) followed by an activity-silent maintenance period. **(B)** IIT predicts consciousness is associated with a maxima of integrated information, Φ, within the “posterior hot zone” (temporo-parietal-occipital cortex) such that that the neural substrate is sustained for the duration of stimulus presentation. **(C)** The dynamic causal modelling (DCM) network architecture used to formally test these predictions. The model comprises the bilateral primary visual cortex (V1), fusiform gyrus (FG; the relevant category-selective region), and the prefrontal cortex (PFC). Green arrows indicate initial sensory input. Black arrows represent extrinsic connections allowed to freely modulate. To evaluate the necessity of specific theoretical mechanisms, targeted connections were “switched off” in reduced models in relevant time-windows. Blue arrows denote the PFC to FG feedback connections removed to test GNWT’s transient broadcasting, while orange arrows denote the FG to V1 feedback removed to test IIT’s sustained posterior connectivity. FG intrinsic self-connections were also tested via Bayesian model reduction and averaging.

In contrast to GNWT, IIT predicts that the physical substrate of a conscious experience (termed the “main complex”) exists as a maxima of irreducible integrated information (Tononi et al., 2016). The most promising candidate network to sustain this complex is argued to likely reside in posterior, sensory regions of the brain – an area that is termed the “posterior hot-zone” (Koch et al., 2016). Temporally speaking, IIT predicts that the neural substrates of consciousness must be actively sustained for the entire duration of the conscious percept (Cogitate Consortium, 2025; Tononi et al., 2016). This is because the theory postulates that the cause-effect structure is identical to the experience itself, so the complex specifying that structure must be continuously maintained

## Current Study & Hypotheses

While the Cogitate Consortium (2025) study was a significant achievement, the original analyses yielded mixed results for both theories. Notably, while some evidence for GNWT onset ignition was observed, the theory’s predicted offset ignition was absent. Further, analyses of inter-regional connectivity challenged both theories. It is possible that the decision to restrict analyses to the high gamma in the original study, may have overlooked potentially critical frequencies within lower frequency bands. To address these possibilities, we re-examined this dataset using DCM for evoked responses to test the connectivity profiles predicted by GNWT and IIT across the whole frequency spectrum.

It is worth noting that the overall design of the experiment was driven by falsificationism (Melloni, 2022); such that all stimuli were clearly visible, fully attended, and there was no additional task to complete and thus, performance was at ceiling levels. Therefore, if GNWT and IIT predict certain brain regions *will* be active in a certain way across time while participants are undeniably conscious of the stimuli, then the predictions of the respective theory should be observed in the data. If these predictions are not observed, then this would pose a challenge to the respective theory.

To test these theories using DCM, we translated the Consortium’s preregistered predictions into specific, diagnostic DCM network predictions (summarized in Figure 2), consisting of two distinct tests each for both GNWT and IIT. While the Cogitate preregistration postulated general relationships between broad cortical zones (e.g., “sustained connectivity between PFC and category-selective areas”), DCM requires a translation of these broad claims into specifically defined network nodes. One of the predictions where this was particularly important was our second IIT prediction. Because IIT predicts that the main complex which instantiates consciousness can move and shrink depending on the exact experience, we assessed whether the postulated sustained connectivity would be instantiated entirely locally (i.e. within the fusiform gyrus; FG). We reasoned that if the FG is embedded within a broader complex -- whether realised locally entirely within the FG or within a broader network between the FG and other (unmodelled) regions – this region would necessarily exhibit sustained self-connectivity. Additionally, we did not test timepoints outside the stimulus onset window for either duration (i.e. 1000ms or 1500ms). Although IIT predicts that the physical substrate of consciousness corresponds directly to the ongoing experience, so that face-related processing must collapse once the face is removed, standard models of visual perception similarly predict a return to baseline activity during a fixation cross as the FFA is highly selective to faces (Kanwisher et al., 1997). Consequently, observing a lack of connectivity at stimulus offset would not uniquely support IIT, insofar as this observation would be similarly consistent with other models of visual processing, rendering the offset period predictions non-diagnostic for IIT.

**Figure 2.**
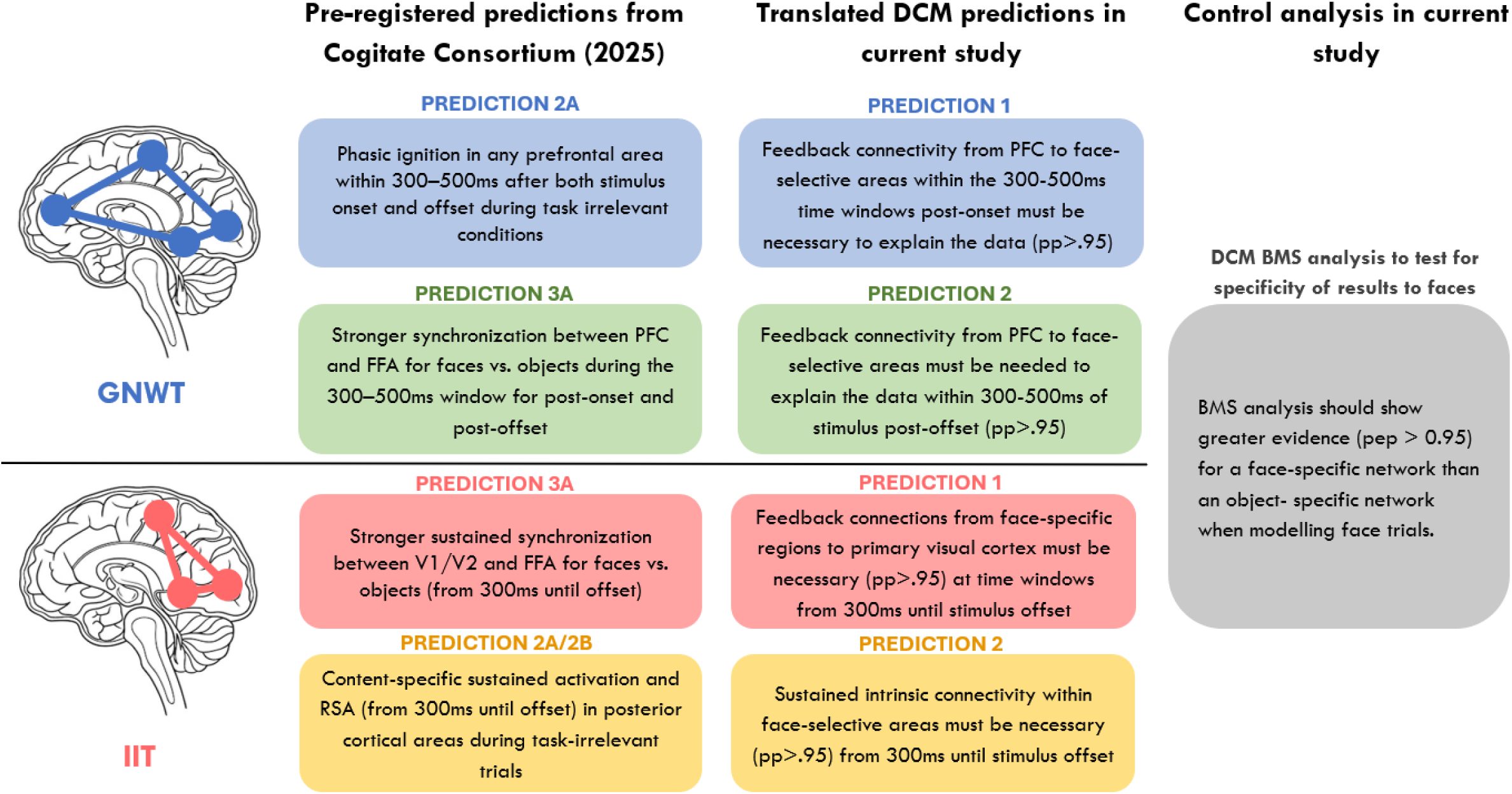
Summary of the key predictions of each theory to our DCM tests. These key predictions were adapted from the preregistered predictions (obtained from Extended Data Table 1) of the original study (Cogitate Consortium, 2025) where the original prediction numbering is referenced. We include only the predictions relevant to our specific analyses, omitting tests of object perception or decoding. The middle column details how these previous predictions were translated into DCM predictions in the current analysis. Note also that for the GNWT predictions we opted to not test the maintenance interval (between onset and offset) because GNWT offers ambiguous predictions during this phase. That is, GNWT predicts either activity-silent maintenance or intermittent bouts of broadcasting depending on whether the stimulus is actively attended (Melloni et al., 2023). The right column describes the control analysis using Bayesian model selection (BMS) which allows one to determine which model best explains the observed data. Here, contrasts of source reconstructed images were used to determine coordinates within regions of interest demonstrating greater activation for objects (i.e. objects > all other stimuli) and faces (i.e. faces > all other stimuli). By comparing models using object-specific versus face-specific coordinates, this analysis was conducted to determine the specificity of the tested networks to face stimuli. This specificity check is particularly critical for evaluating IIT, as this theory makes strong claims regarding the content-specificity of the underlying neural substrates. Posterior probability is denoted by pp, with a threshold of pp > 0.95 indicating strong evidence for the model (Kass & Raftery, 1995).

To test these predictions, we focused on a specific subset of the Cogitate Consortium’s experimental data. To isolate the neural correlates of conscious perception and reduce task-related executive or motor confounds, we restricted our analysis entirely to task-irrelevant trials. Additionally, we specifically analysed face stimuli because they reliably engage a well-characterised higher-sensory network, allowing us to anchor our models to the FG; a well-established higher-sensory, category-selective area (Kanwisher et al., 1997), that is also argued to be a content-specific region for conscious face perception (Koch et al., 2016). We also constrained our analyses to the 1s and 1.5s stimulus durations to ensure a sufficiently long temporal window to evaluate IIT’s prediction of sustained connectivity. Additionally, because of the well-known feedforward sweep of information across the cortical hierarchy which accompanies visual perception, it is only the feedback connections that are able to theoretically distinguish these theories of consciousness from standard models of visual perception, and thus we constrained our analysis to testing only the feedback connections. Finally, we opted to only analyse the magnetoencephalography (MEG) dataset here (not the fMRI/EEG). MEG offers both the millisecond-level temporal resolution needed to test precise predictions about the timing of neural responses and also offers spatial resolution advantages needed for cortical source reconstruction.

As the differing predictions diverge not only spatially, but also temporally, we used an expanding time-window approach for our DCM inversions (inspired by a similar approach by Garrido et al., 2007). We modelled data across expanding time windows following both stimulus onset and offset, to track model evidence over time and identify when specific directed connections emerged and whether they were sustained throughout the duration of the percept. Finally, to ensure the robustness and replicability of our findings, we adopted a split discovery and validation procedure (Poldrack et al., 2017).

## Methods

Data for this experiment were collected by the Cogitate Consortium (2025) and kindly made publicly available. For the sake of context, a brief overview of the experiment design will be provided here but for detailed methods regarding data acquisition and participant demographics, please refer to the original paper (Cogitate Consortium, 2025).

The entire analysis pipeline, including preprocessing, source reconstruction, DCM specification, and PEB, was developed and finalised exclusively on the discovery batch. Once the pipeline was fixed, it was applied without modification to the validation batch. Therefore, in addition to the predictions of either theory being tested here, the splitting of analyses required that the predictions be present in both batches of analyses to be considered robust and replicable. Significant findings in one batch but not the other must be interpreted with caution given their lack of replicability.

### Participants

A total of 100 healthy volunteers were recruited for the MEG portion of the study. Data was split into two batches for analysis. This included a discovery set of N = 48 participants and a validation set of N = 52 participants. There was 1 exclusion from batch 1 and 2 because they failed to achieve the required hit rate of >80% or a false alarm rate higher than 20%. This left a total sample of 98 participants.

### Design and Procedure

As illustrated in Figure 3, participants viewed a sequence of suprathreshold visual stimuli while performing a non-speeded target detection task. Stimuli consisted of four categories: faces, objects, letters, and false fonts; each of which were comprised of a set of 20 different identities (e.g. 20 different face identities). These were presented for three different durations (0.5s, 1s, and 1.5s) to manipulate the length of the conscious percept and all stimulus presentations plus the subsequent blank screen were fixed at a length of 2s. This ensured a minimum post-stimulus window of 0.5s to capture stimulus-offset responses prior to the onset of the next stimulus. In each block, participants were instructed to detect two specific target identities (e.g., a specific face and a specific object out of the 20 possible identities) among a series of stimuli. This effectively divided the stimulus stream into three distinct classes of stimulus type. First, targets, were the specific identities which required a response. This was to ensure participants were attending to and completing the task. The second type were task-relevant non-targets, which were stimuli belonging to the same category as the targets (e.g. other face identities) but not matching the specific target identity in a given block. These stimuli were therefore relevant to the task at hand but did not require a motor response. The final type, task-irrelevant, were stimuli which did not belong to the same category as the targets. These were consciously perceived (i.e. suprathreshold) but were irrelevant to the current task. By counterbalancing the target categories across blocks (e.g., detecting faces or objects in one block, and letters or false fonts in the next), the design ensured that every stimulus category served as both relevant and irrelevant throughout the experiment.

**Figure 3.**
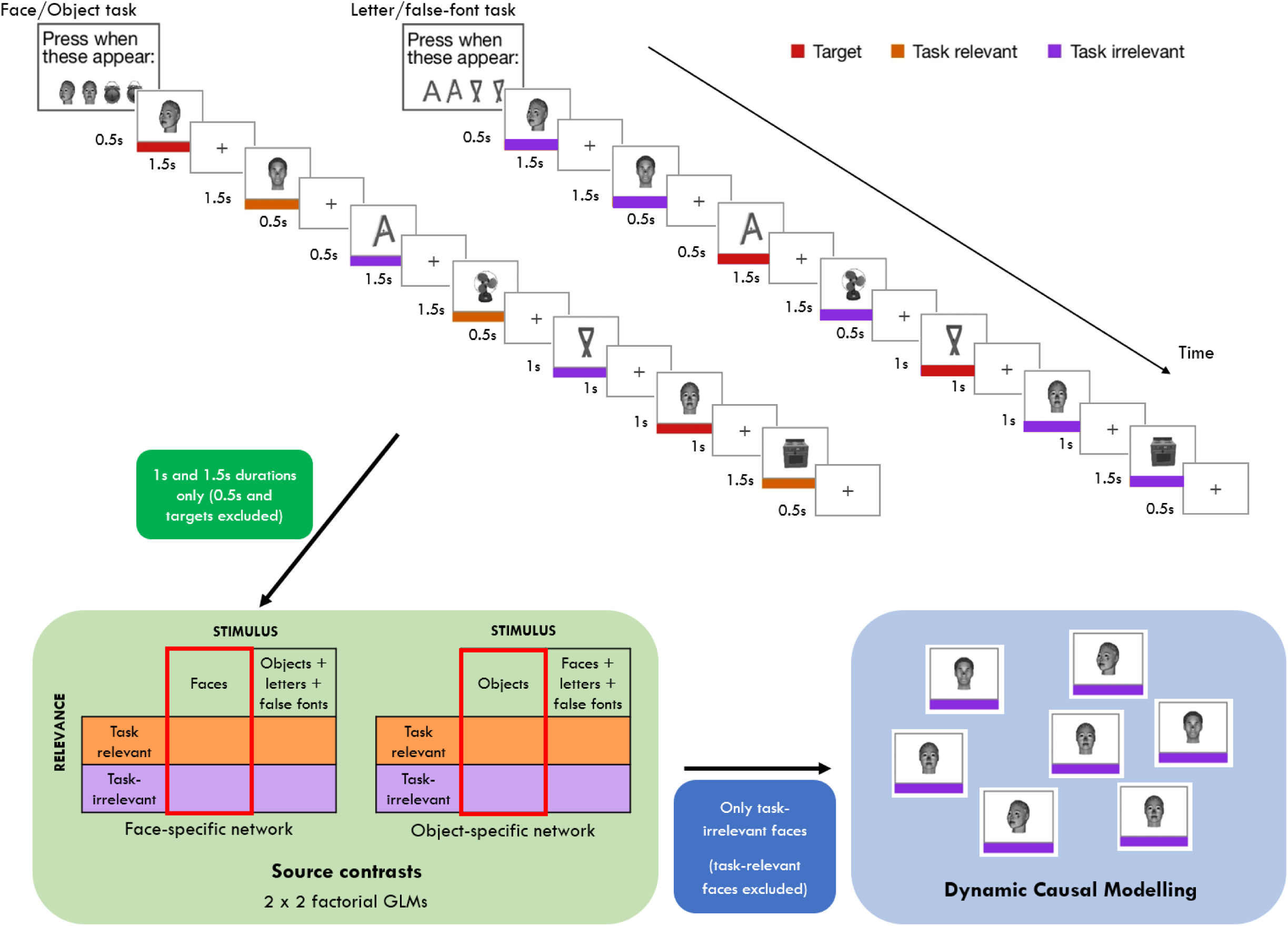
Schematic of the experimental design adapted from the Cogitate Consortium (2025) and the key trials selected in each step of the current analysis. Participants viewed a stream of suprathreshold stimuli from four categories (faces, objects, letters, and false fonts) presented for varying durations (0.5, 1.0, or 1.5 s) followed by a blank screen (2s minus stimulus duration) such that each stimulus and blank screen was presented for a total of 2s. The experiment employed a block design to manipulate task relevance. In “Face/object” blocks (left stream), participants were instructed to detect specific target identities (e.g., a specific face and object). Stimuli were classified into three conditions: targets (red bars; target identities requiring a response), task-relevant (orange bars; non-target stimuli belonging to the target category, e.g., other face/object identities), and task-irrelevant (purple bars; stimuli from non-target categories, e.g., letters/false-fonts). In “Letter/false-font” blocks, the target categories were reversed, rendering letters and false fonts task-relevant while faces and objects became task-irrelevant. Note also that the coloured bars were not presented in the actual experiment and are only presented here to illustrate the different stimulus types. The bottom section shows the trials that made it to the main analysis in the current study, where first target stimuli and any 0.5s duration stimuli were excluded and all other trials with the face-specific GLM used to identify a face-specific network and a second GLM to identify an object-specific network, used for the control analysis. The red square denotes the main effect comparison made to identify each stimulus specific network. From here, DCMs were inverted on only the task-irrelevant face trials for both face-specific and object-specific networks.

### MEG preprocessing

Raw MEG data, acquired from 306 sensors (102 magnetometers, 204 planar gradiometers) was preprocessed using a combination of MNE-Python (v1.8.0; Gramfort et al., 2014) and SPM12 (https://www.fil.ion.ucl.ac.uk/spm/) in MATLAB 2024b (The MathWorks Inc., Natick, MA). All data was pre-processed and analysed on the high-performance computing platform, Spartan, at the University of Melbourne. This pipeline leveraged the strengths of MNE-Python for signal denoising (specifically, Maxwell filtering and FastICA) with SPM12 for its native support of DCM and Bayesian source reconstruction. The raw MEG data underwent Maxwell filtering (Signal-Space Separation; SSS; Taulu & Kajola, 2005) to suppress environmental noise and sensor artefacts. Prior to SSS, noisy and flat channels were automatically detected using the “find_bad_channels_maxwell” function, with the aid of cross-talk compensation and calibration files. These marked bad channels were reconstructed during SSS.

To remove artefactual sources in the data, ICA was computed on a separate copy of the continuous sensor data after it was bandpass filtered from 1 to 40 Hz and downsampled to 200Hz. Here, the FastICA algorithm (Hyvarinen, 1999) was used to extract as many components as necessary to explain 99% of the variance within each participant. Artefactual components, primarily those related to eye blinks and cardiac activity, were identified and then rejected via visual inspection of their topography and time course. An average of 3.19 components (SD = 0.98 components) were removed per participant across both batches.

The cleaned continuous data was then converted to SPM readable format and the remaining preprocessing steps were completed in MATLAB. Bad segments were marked where channels exceeding a threshold of 5000fT or planar gradiometer channels exceeding 5000fT/cm were marked as artefactual. Trials containing such artifacts, or where more than 20% of channels were bad, were subsequently marked for rejection. On average, 103 trials made it to the analysis stage for each participant.

To evaluate the distinct temporal predictions of the theories, the continuous data were branched into two separate epoching pipelines: one aligned to stimulus onset, and a second aligned to stimulus offset.

For the onset aligned epochs, data from all stimulus categories were segmented into epochs spanning -100ms to 2000ms relative to stimulus onset. Retaining both task-relevant and irrelevant trial types at this stage was necessary to allow for the subsequent GLM source contrasts (i.e., face trials versus all other stimulus types). However, because we were specifically interested in sustained neural responses, trials with a 500ms stimulus duration were excluded; only the 1000ms and 1500ms duration trials were retained for analysis.

To test the GNWT prediction of an offset ignition, a second set of epochs were created for the 1000ms and 1500ms trials. Here, we were first re-aligned these trials to stimulus offset (defined as 0ms) and segmented into epochs spanning -100ms to 600ms relative to offset.

For both the onset and offset aligned epochs, all trials within each condition using the robust averaging procedure. This procedure down-weights outlier time points within trials before averaging. To remove any high-frequency noise potentially introduced by robust averaging, the averaged data were low-pass filtered at 40 Hz (Litvak et al., 2011). Finally, baseline correction was applied using an interval of -100ms to 0ms relative to stimulus onset or offset. While this baseline correction step for the offset-aligned epochs may not be a neutral resting state (i.e. the time window being baseline corrected is during face perception), baseline correcting with respect to the offset was chosen to isolate the neural dynamics evoked by the stimulus removal (i.e., during the updating of a percept at offset predicted by GNWT).

### Identifying regions of interest for the DCM analysis

Source reconstruction was conducted in SPM. Individual structural MRI scans were used in the inversion procedure to form a single shell model for each participant’s head mesh during the source reconstruction. In SPM, these individual meshes are generated by mapping and transforming their structural image against a canonical template brain, allowing for more robust solutions when creating the head meshes (Mattout et al., 2007; Ashburner et al., 2014). For those without a structural MRI scan (a total of five participants), a template mesh was warped to the participant’s fiducials. Each participants’ cortical mesh had 8196 vertices.

For the onset-locked data, these models were subsequently inverted using a greedy search algorithm across a time window of 0 to 2000ms relative to stimulus onset, and a frequency window spanning from 0 to 256 Hz to capture all relevant cortical frequencies. The greedy search algorithm is a variant of the multiple sparse priors approach within SPM. To ensure a consistent spatial model across both onset and offset effective connectivity analyses, the inverse solution derived from this broad 0-2000ms window was subsequently used for the offset-locked epochs. As this inversion was used only to define a consistent spatial model for subsequent DCM analyses rather than to characterise temporal dynamics (which are modelled in the DCM analyses) the time window used for source reconstruction was not constrained to the duration of the stimulus presentation.

These source estimates were converted to images for subsequent analysis, and we focused on capturing the peak sources of activation during the onset of the face stimulus within each region of interest (ROI). As such, the images used in the subsequent GLM analyses were created by projecting the source estimates onto a standard MNI mesh during a time window of 0 to 600ms relative to stimulus onset. The resulting images were smoothed to 12mm^3^.

To adjudicate between GNWT and IIT, we required a network architecture capable of testing both posterior network processing and global broadcasting. Therefore, our models comprised three bilateral cortical levels. First, we required a lower visual node to receive the initial afferent visual input (primary visual cortex; V1). Second, we included a higher-level category-selective node relating to brain areas known to be involved in processing of faces (FG). Anchoring the network around the FG was critical, as both theories make explicit, competing predictions regarding the temporal and directed connectivity of this specific node during face perception. Finally, we included a higher-order cognitive node to test for global workspace broadcasting (PFC).

To define the coordinates for each node within the DCM architecture, we took a hybrid approach driven by both the individual’s data and the key hypotheses of interest. A 2 x 2 full factorial generalised linear model (GLM) was fitted to the source reconstructed images for each participant separately. The 2 factors in the model were stimuli (face or all other stimuli; i.e. objects, letters, and false fonts) and task-relevance (relevant or irrelevant). Both factors’ levels were assumed to be dependent and have equal variance. For each ROI, an anatomical mask was created separately for each hemisphere using JuBrain Anatomy Toolbox (Eickhoff et al., 2005). To create the frontal mask, all frontal regions were selected (Amunts et al., 1999; Amunts et al., 2004; Bludau et al., 2014; Henssen et al., 2016; Geyer, 2012; Geyer et al., 1996). For the fusiform mask, all fusiform gyrus regions were selected (Caspers et al., 2013; Lorenz et al., 2017). And to create the V1 mask, the V1 region was selected (Amunts et al., 2000). For each contrast we analysed the main effect of face stimuli (i.e. faces > all other stimuli) and the peak coordinate was then extracted for each ROI constrained over the corresponding anatomical mask for each participant. This allowed us to optimise the specific MNI coordinate to the most robust voxel of activation elicited by faces for each participant, at each ROI. Additionally, we determined the regions demonstrating greater activation for objects compared to all other stimulus categories (i.e. objects > all other stimuli). Models with these ROI coordinates were then compared against models using the face-specific ROI coordinates using Bayesian model selection (which we cover in more detail below). This analysis was conducted to test for the specificity of the face processing network; such that a network with peak activation for face stimuli should explain face stimuli better than a network whose peak activation occurs for objects.

The custom-fitting of the DCM network meant that, for each participant, the specific peak MNI coordinate fed into the DCM model was functionally localised for this contrast of interest. This procedure was especially important for the PFC nodes. GNWT considers the PFC broadly relevant and remains largely impartial to the subregions within the PFC which are specifically important. Further, as the PFC is quite large and we have previously found that PFC activation does not overlap much between participants (Bandara et al., 2025), this meant the group-level PFC coordinate may not be representative of each participant. Thus, this allowed us to best capture any influence from the PFC should they exist.

### Effective connectivity analysis

Effective connectivity was estimated using DCM for evoked responses (David et al., 2006) specified as an Equivalent Current Dipole (ECD) spatial model using a Jansen and Rit neural mass model (Jansen & Rit, 1995). Only task-irrelevant faces were modelled here to minimise confounds of motor preparation for reporting.

As the temporal evolution of connectivity was an important empirical question here, we opted for an expanding time-window approach (similar to Garrido et al., 2007). Here, separate DCMs were inverted for each participant in expanding time windows from 0ms to 1000ms post-stimulus onset with 100ms increments (e.g. 0ms to 100ms, 0ms to 200ms, 0ms to 300ms, etc.). The rationale of this expanding time-window approach was to track the model evidence across these time windows to identify when specific connections became necessary to explain the data, and if these remained necessary throughout the duration of the stimulus presentation. Similarly, to test for offset responses, we ran another set of DCMs starting from 0ms to 300ms after stimulus offset and continuing up to 500ms in steps of 100ms again (i.e. 0ms to 300ms, 0ms to 400ms, and 0ms to 500ms post-stimulus offset). Note that for these offset DCMs, the afferent input volley was specified exactly the same as the onset. That is, the input was specified over the early sensory nodes using a prior of 64ms and allowing the model to determine the precise input timing and duration.

### Hierarchical Bayesian modelling

Once the expanding time-window DCMs were inverted for every participant, parametric empirical Bayesian (PEB) modelling was used to investigate the strength of the connectivity parameters at the group-level. This form of modelling involves conducting a second-order hierarchical Bayesian model over the connectivity parameters across the group (Friston et al., 2016). As only one condition was modelled in this case, the PEB was inverted over the A-matrix. Additionally, as we modelled the DCMs across multiple time windows, we conducted one PEB on all participants per time window.

To arbitrate between the theories, Bayesian model comparison was used to compare the fully connected model against reduced models where specific connections important to either theory were “switched off”. Switching off or removing connections is equivalent to fixing these connections at a prior mean of zero in log-scaling space, corresponding to their canonical default strength (in this case, 16Hz). As computing the evidence for a given model, *P*(*y*|*m*), is computationally intractable, this is approximated within the PEB framework as free energy. Further, as the free energies across different time windows are not comparable because they are derived from by different data (Stephan et al., 2010), the difference in free energy was computed between the full and the reduced models within each time window. This difference was then converted to a posterior probability for the full model compared to the reduced model using the equation:

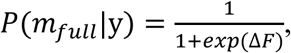

where Δ*F* represents the log Bayes factor in favor of the reduced model (that is, F_reduced_ − F_full_).

### Theoretical translation and model specification

To test the key predictions of GNWT, that conscious perception relies on a transient prefrontal broadcasting at stimulus onset and offset, we constructed a reduced model where bilateral backward connections from the PFC to the FG were removed. If the full model (with PFC feedback) had meaningfully higher free energy compared to the reduced model, it would support the GNWT prediction that PFC feedback is necessary for conscious perception. That is to say, this would suggest that removing PFC feedback connections impacts the model’s explanatory power, and hence those connections are important to explain the data. As GNWT predicts this response should occur within 300-500ms of stimulus onset (prediction 1) or offset (prediction 2), we compared these reduced PEBs to the full model for each of these time windows separately and then computed their posterior probability.

To test the predictions of IIT, that conscious perception requires a sustained content-specific neural response tracking the duration of the percept, we constructed a reduced model by switching off key connections within a posterior content specific network – specifically, backward connections from the FG to V1 (prediction 1). Again, evidence favouring the full model over this reduced version, would support IIT’s prediction of posterior integration. Because IIT hypothesises posterior integration should be sustained for the duration of the perception (from 300ms onwards), the PEBs were compared for each time point starting from 0ms to 300ms and up until 1000ms relative to stimulus onset.

As a further test of IIT, we investigated whether the sustained response in the ‘posterior hot zone’ might manifest as sustained self-connectivity within the content specific region (i.e. the FG). To do so, we analysed the G-matrix of the DCM, specifically, its excitatory receptor density parameters (*H*e). While these parameters are not explicitly a connectivity parameter, as Kiebel et al. (2007) have demonstrated, they behave like one; as they increase the overall influence the subpopulations within a cortical column have on each other, such that changes in these parameters modulate the magnitude of the overall activity within an area/node.

Consequently, these parameters serve as a suitable proxy to the node’s intrinsic connectivity. For this test, we performed an automated PEB search to track the posterior probability that intrinsic connections in the FG were needed to explain the data. This process involves performing Bayesian model reduction to search across a vast model space by switching off connections in the model iteratively, comparing these models to a fully connected model, and then pruning redundant connections which do not positively contribute to the overall model evidence. Bayesian model averaging (BMA) is then performed over the parameters of the final set of models to yield an estimate of the group-level connectivity parameters. We then traced the posterior probability of these connections over time from 0ms to 100 to 1000ms within the FG.

In the context of DCM, where local (intrinsic) dynamics are parameterized by the excitatory post-synaptic gain of a node, any such sustained activity that cannot be explained by extrinsic parameters would be mathematically absorbed by this intrinsic parameter (Friston et al., 2003; Kiebel et al., 2007). Therefore, we can treat the self-connectivity FG parameter as a proxy for the region’s specific involvement in a self-sustained complex – irrespective of whether that complex is localised within the FG or a larger network. In this context, sustained evidence for intrinsic connectivity in the FG would support the hypothesis that this region is involved in a sustained network during stimulus viewing.

All results here are reported as the posterior probability of the connection being present using a threshold of pp > 0.95, which constitutes strong evidence for the necessity of the connections to explain the data (Kass & Raftery, 1995).

### Bayesian Model Selection

As a final test, we evaluated content specificity of the DCM network to face processing post-hoc. To do so, a validation analysis was run to compare the face-specific network against object processing. Here, identical DCMs were run using an object-specific network (i.e. taking the peak MNI coordinates when contrasting objects against all other stimulus categories). These DCMs were inverted across the same expanding time windows (from 0–100ms up to 0–1000ms). Random-effects (RFX) BMS (Stephan et al., 2009) was then employed to compare the models at the group level. For each time window, we computed the protected exceedance probability (PEP), which measures the probability that a given model is more likely than any other model tested, over and above chance (Rigoux et al., 2014). This provided a test of whether the face-specific network provided a more parsimonious explanation of the empirical data than the generic object network, suggesting that the DCM network was specific to faces. It is worthwhile noting that these analyses were conducted after the primary hypothesis-driven DCM inversions were complete and is reported here in the order in which it was conducted.

## Results

### Nodes identified for DCM analysis

We extracted the peak MNI coordinate for each participant resulting from the main effect of face stimuli, for each node to account for individual variability. This node optimization process was particularly important for the PFC nodes, where prefrontal activation exhibited a high level of between-subject variability (see Table 1). To summarise the spatial location of this network, we computed the mean and spread (standard deviation) of the individual peak coordinates determined in the source reconstruction procedure. As shown in Figure 4, these group-average coordinates represent the spatial centre for each of the modelled ROIs. Furthermore, the size of each spherical node reflects the between-subject spatial variability at that source, derived from the mean standard deviation across the x, y, and z dimensions. The resulting group-average coordinates across both datasets corresponded to bilateral early visual cortex (V1/V2; BA17/18), the fusiform gyrus (Fusiform Face Area; BA37), and the inferior frontal gyrus (PFC; BA45). The source inversion yielded a mean variance explained for the source solution of 77.36% (SD = 5.05%) for the discovery batch and 76.88% (SD = 3.90%) for the validation batch, indicating a robust fit of the forward model to the sensor data. One participant in the validation batch failed to show any suprathreshold activation in the left PFC node. As such, for this participant, the group-level (specifically, at the batch-level) peak coordinate was used for that node.

**Figure 4.**
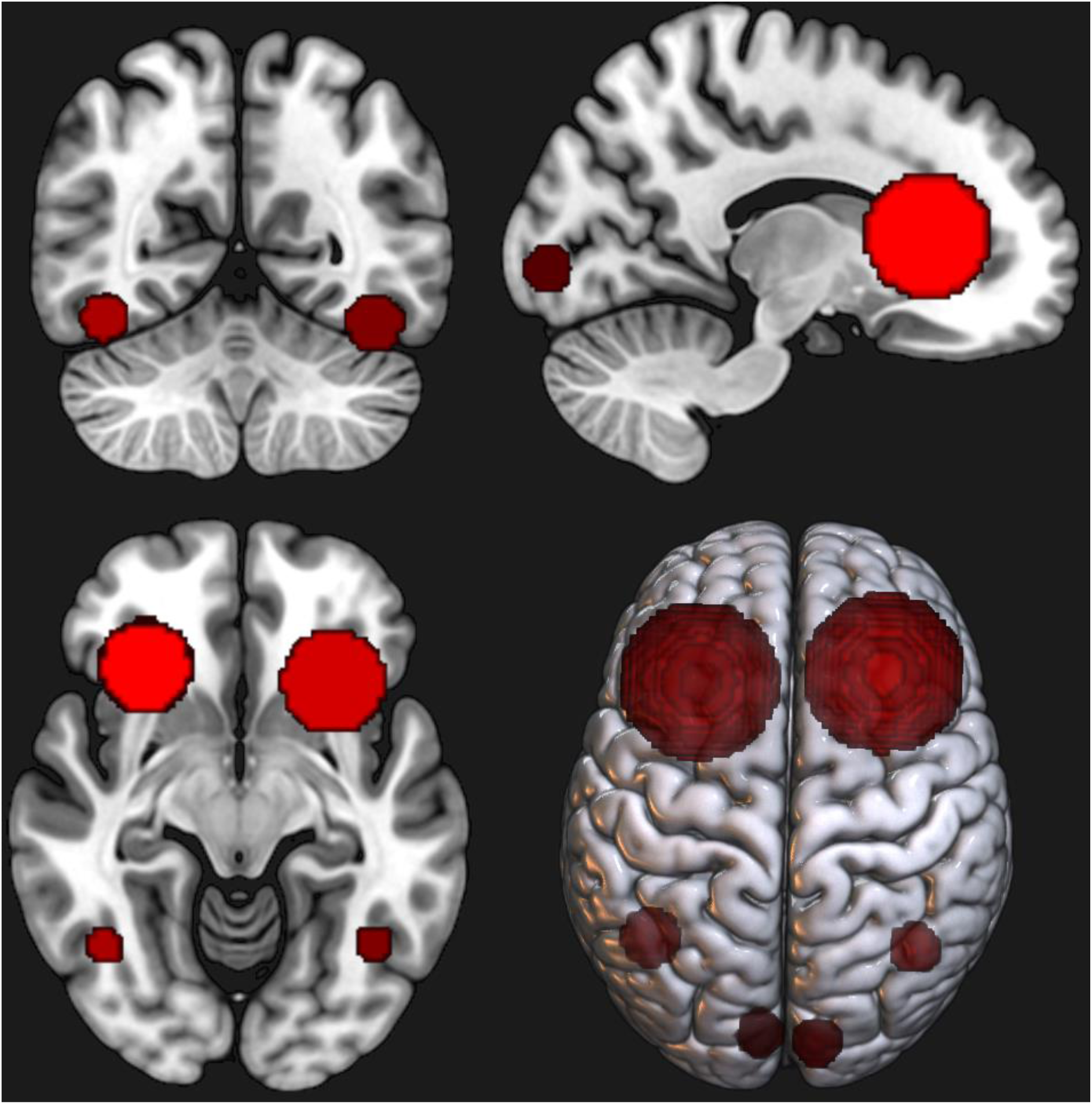
Group-average source locations across both batches for the six-node DCM network. Nodes are centred at the mean peak MNI coordinates across all participants across both batches, derived from individual subject-level GLM contrasts (faces > objects) within anatomical ROI masks. Sphere size reflects the spatial variability of peak locations across participants, derived from the mean standard deviation across the x, y, and z dimensions at each source.

**Table 1.** Group-average MNI coordinates and corresponding Brodmann Area. Coordinates represent the mean peak MNI location across participants. SD denotes the standard deviation of the distance from the mean coordinate for each axis.

| Region of Interest | Brodmann Area | *x* (SD) | *y* (SD) | *z* (SD) |
| --- | --- | --- | --- | --- |
| L Early Visual (V1) | BA 18 | -6.5 (5.5) | -89.1 (6.9) | -0.6 (10.4) |
| R Early Visual (V1) | BA 17 | 12.0 (7.2) | -91.4 (7.9) | -0.1 (7.5) |
| L Fusiform (FG) | BA 37 | -41.9 (6.0) | -59.8 (15.0) | -16.7 (5.2) |
| R Fusiform (FG) | BA 37 | 41.5 (4.8) | -60.6 (12.5) | -15.1 (5.0) |
| L Prefrontal (PFC) | BA 45 | -29.3 (15.6) | 21.7 (24.0) | 8.6 (33.1) |
| R Prefrontal (PFC) | BA 45 | 29.0 (17.2) | 26.3 (27.2) | 10.4 (29.0) |

### Testing PFC to FG connectivity at onset and offset

We focused our analyses on testing only the specific connections required to evaluate the predictions of GNWT and IIT. To demonstrate replicability, we report the findings for the discovery batch first, followed immediately by the corresponding results from the independent validation batch for each prediction.

To test GNWT’s first prediction, we used Bayesian model comparison (BMC) to compare a fully connected model against a reduced model where bilateral backward connections from the PFC to the FG were “switched off” (fixed at a prior mean of zero in log-space; see Figure 5a). In our initial discovery stage analysis, we observed strong evidence (pp > 0.95) for the necessity of key GNWT connections during the 300–500ms window post-stimulus onset. These results were replicated in the validation batch (pp > .95).

**Figure 5.**
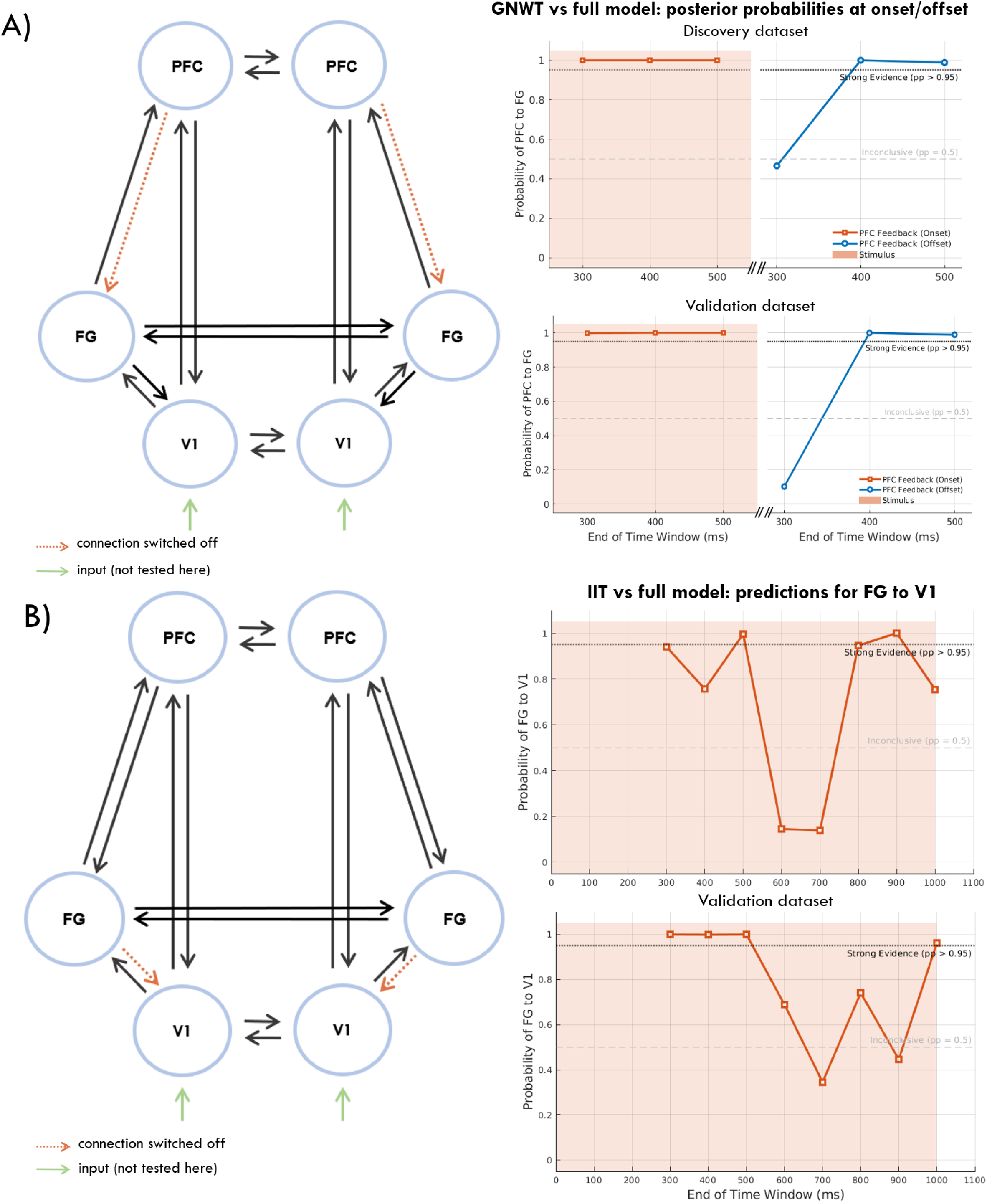
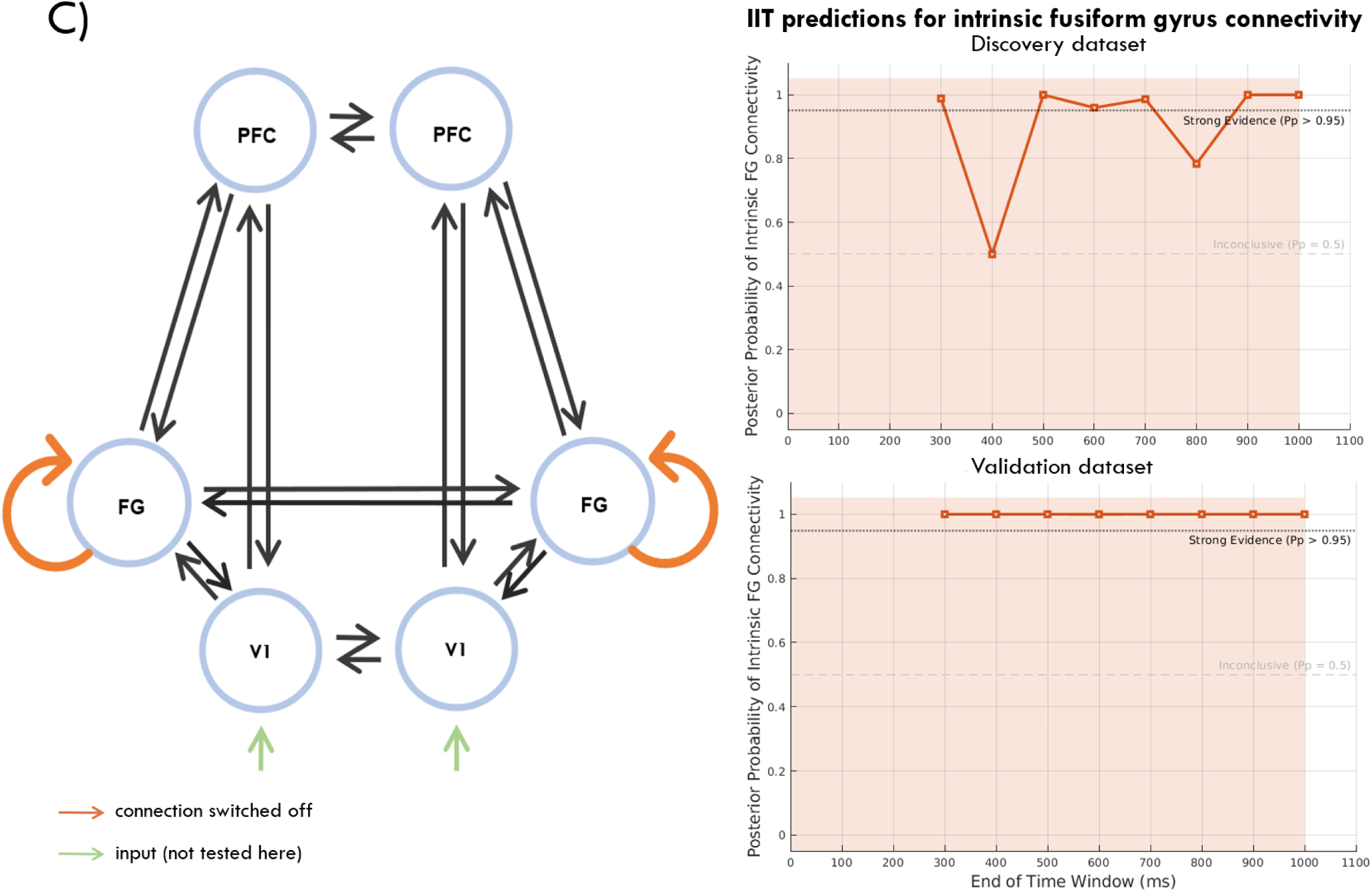
Expanding Time-Window Bayesian Model Comparison of Reduced DCMs Tests of GNWT and IIT. Each panel assesses the necessity of specific connections predicted by GNWT or IIT. Here, evidence for the full model is compared against reduced models where those connections were “switched off” (log-scaling parameters were fixed at zero). On the left of each panel is a schematic illustrating the set of connections being evaluated (highlighted in orange). On the right are line plots showing the time course of the posterior probability (pp) that these specific connections are necessary to explain the data. In each panel, the top graph shows the discovery dataset and the bottom graph shows the independent validation dataset. **A)** Test of GNWT’s predictions 1 and 2 of prefrontal ignition at stimulus onset and offset. The analysis evaluates the necessity of long-range top-down feedback from PFC to FG. Separate models were run for stimulus onset (orange lines) and stimulus offset (blue lines) to test for “ignition” events. **B)** Test of IIT prediction 1. The analysis evaluated the necessity of feedback between V1 and FG. Sustained high probability (pp > 0.95) indicates strong evidence for the putative posterior network connections being necessary to explain the data for each time step. **C)** Test of IIT prediction 2. This analysis examined sustained intrinsic connectivity within FG. The analysis evaluates the necessity of the excitatory gain parameter (a proxy for intrinsic connectivity within the node; Kiebel et al., 2007) within the fusiform gyrus (FG) nodes at each time step. In all plots, the y-axis represents the posterior probability that the relevant connections/parameters are necessary for model fit. The dotted black line at pp = 0.95 indicates the threshold for strong evidence. The shaded orange background indicates the period where the visual stimulus was present on screen.

Equivalently, we ran the same set of analyses at stimulus offset to test GNWT’s second prediction. Again, we compared a full model against a reduced one without feedback connections from the PFC to the FG. Across both the validation and discovery datasets, we observed strong evidence (pp > 0.95) for the necessity of the full model containing key GNWT connections, however starting from 400ms window post-stimulus onset (pp > .95). These findings are in line with the GNWT prediction of a phasic “ignition” event at offset.

### Testing sustained FG connectivity

To test IIT’s first prediction, we again used BMC to compare a fully connected model against a reduced model where the key connections IIT deems necessary for conscious face perception were switched off (i.e. feedback FG to V1 connections; see Figure 5b). In the discovery batch, feedback from FG to V1 did not show a sustained response throughout the stimulus duration, with the posterior probability of the connections only being necessary to explain the data (pp > 0.95) for only two (non-successive) time windows out of the eight time-windows tested. Similarly, in the validation dataset we did not observe a sustained response from the FG to V1 throughout stimulus presentation. While evidence for these backward connections was initially robust (pp > 0.95 between 300ms and 500ms), the posterior probability subsequently dropped, fluctuating around the inconclusive threshold (pp ≈ 0.50) for much of the later time windows, before unexpectedly rebounding to strong evidence in the final 1000ms window.

Notably, in our tests we observed substantial volatility in the posterior probabilities (e.g., the sharp drops in evidence at time windows ending at 600ms and 700ms followed by a rebound at later windows in the discovery batch). We sought to rule out that there might have been inversion failures in the underlying DCMs; specifically that certain time windows may have been stuck in local (c.f. global) optima of free energy. Because each time window is inverted independently, a poor inversion at one window (but not its neighbours) could in principle produce a spurious drop in group-level evidence that reflects an optimisation failure rather than a bona fide change in coupling. To test this possibility, we examined the mean first-level free energy across the expanding time windows. Because free energy scales with the amount of data being modelled, and each successive window incorporates more peristimulus samples, the mean free energy would be expected to decline smoothly and monotonically as the window lengthens. If the observed volatility in posterior probability was indeed driven by inversion failures, it would be expected that these would coincide with corresponding spikes in mean free energy in the DCMs for those time windows. Our assessments confirmed that the mean free energy (an approximation of model evidence) behaved as expected for both discovery and validation batches, indicating a stable model convergence (that is, the dips in the group-level posterior probabilities did not correspond to dips in free energy at the first-level DCMs; see Figure 7).

**Figure 6.**
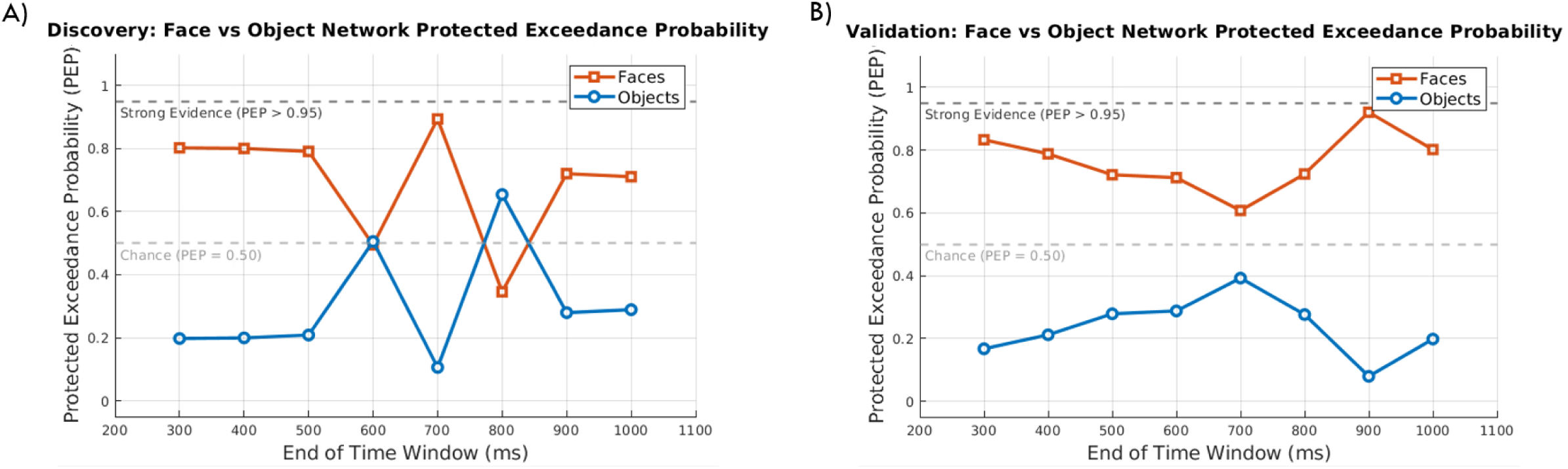
Bayesian model selection comparing face-specific and object-specific networks when modelling face trials. Protected exceedance probability (PEP) for the discovery **(A)** and validation **(B)** dataset across expanding time windows. The face-specific network (orange squares) generally provides a more parsimonious fit than the object-specific network (blue circles), remaining mostly above chance (PEP = 0.50). However, it does not surpass the threshold for strong evidence (PEP > 0.95) at any time window. Overall, while data trends positively towards face-specificity, the absence of strong evidence indicates that these content-specific connectivity differences should be interpreted with caution, as the data may not be well-suited to draw strong conclusions.

**Figure 7.**
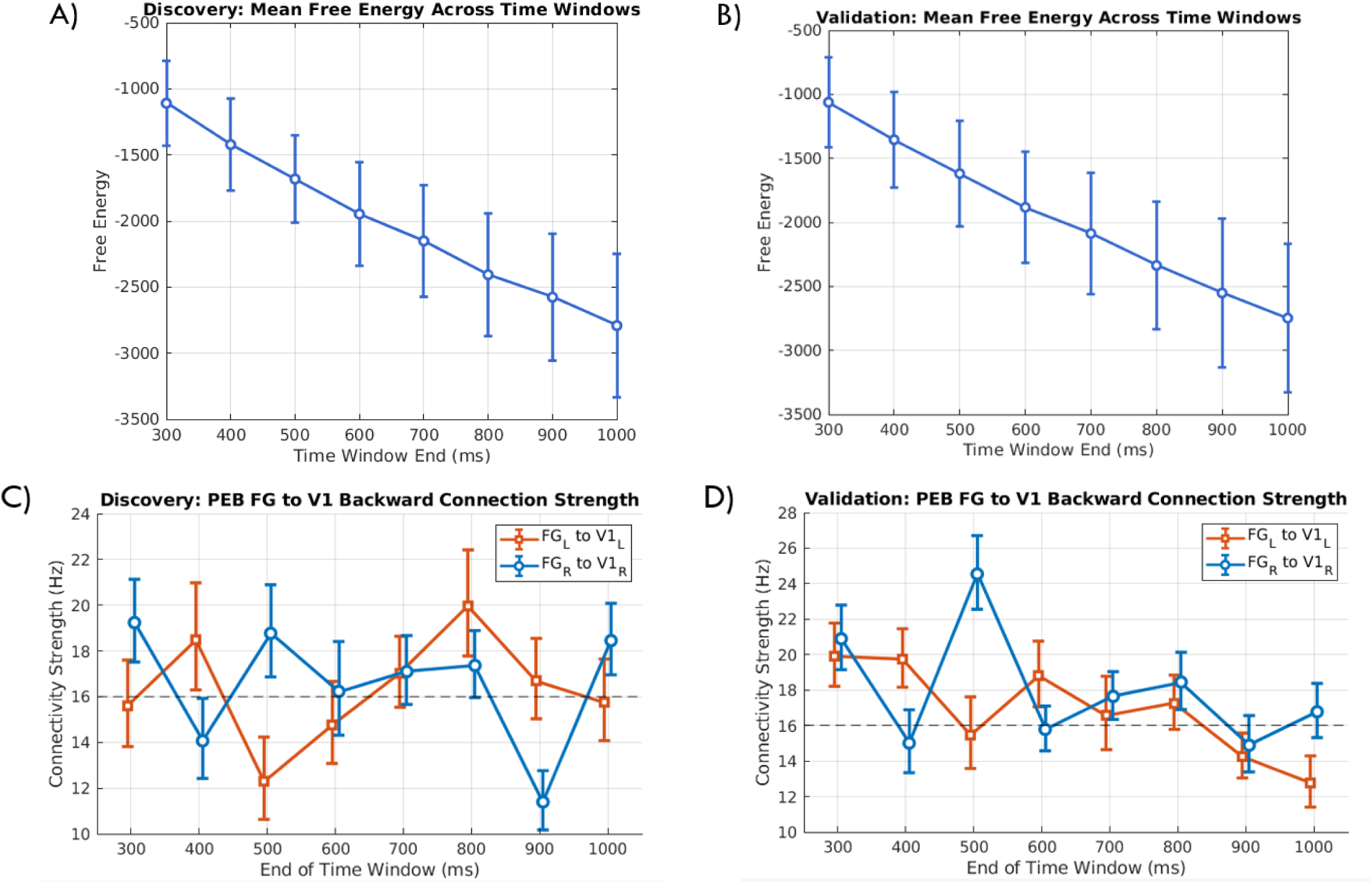
First-level convergence diagnostics and group-level Parametric Empirical Bayes (PEB) parameter estimates across expanding time windows. Here, in the discovery (**A**) and validation (**B**) datasets, we highlight the gradual decrease in mean free energy across all participants demonstrates expected model behaviour as the time window length increases (A and B; free energy is related to, among other things, the number of samples being modelled), and a lack of a substantial dip at the corresponding time windows where posterior probabilities dip confirms the absence of gradient ascent inversion failures (i.e. local maxima problem). Having established this, we then plot the group-mean effective connectivity parameters for the IIT feedback connections tested (fusiform gyrus [FG] to primary visual cortex [V1]) in the discovery (**C**) and validation (**D**) datasets. The transient dip of these estimates towards the prior value of 16Hz (dashed line) coincides with the observed drops in posterior probability for the full model as seen in the corresponding test of IIT (prediction 1; i.e. at time windows ending at 600ms and 700ms in the discovery batch). Note, in A and B, error bars indicate between-subject standard deviation. In, C and D, blue lines represent right-hemispheric connections and orange lines represent left-hemispheric connections. Error bars denote 90% Bayesian credibility intervals.

We subsequently examined the temporal evolution of the group-level connectivity strengths against the main results for the first prediction (Figure 7C and D). This inspection revealed that the group-level connectivity strengths (and their credibility intervals) shrank towards zero during time windows where the posterior probabilities dropped. Subsequently, an inspection of the connectivity parameters at some later time windows revealed that a late-stage rebound in posterior probability was primarily driven by a strong decrease in connectivity.

For IIT’s second prediction, we evaluated intrinsic connectivity via the G-matrix in DCM (see Figure 5c). In the discovery set, there was evidence for sustained intrinsic connectivity within the FG from 300ms onwards. Although the posterior probability of the parameter being needed to explain the data fell to an inconclusive level at the 600ms time window, it rebounded in subsequent windows returning to above the 95% threshold for the remainder of the epoch and finishing with positive evidence (pp ≈ 0.78; Kass & Raftery, 1995). This behaviour was not replicated in the in the validation dataset however, where we observed the necessity of this parameter sustained in all time windows for the entire stimulus duration (pp >0.95).

### Bayesian model selection control analysis

Following the testing of each theory’s predictions, we sought to empirically validate the content-specificity of our network architecture. RFX-BMS was used to confirm whether the face-specific network provided a more parsimonious explanation of the data than an object-specific network when modelling face trials. For each time window, the PEP was computed to determine which model provided a more parsimonious account of the data, over and above chance. For both the discovery and validation batches, the PEP across the expanding time windows revealed a trend favouring the face-specific network but did not reach the threshold for strong evidence at any time window in either dataset (see Figure 6).

## Discussion

In this study, we re-analysed the Cogitate Consortium’s (2025) MEG dataset using effective connectivity methods to adjudicate between two popular theories of consciousness, IIT and GNWT. Across four different predictions, we found results consistent with both GNWT predictions, while IIT’s first prediction was challenged and somewhat consistent on the second prediction. Despite results for these predictions broadly being replicated across both batches of analysis, a subsequent validation analysis comparing a face-specific network to an object-specific network to test for the content-specificity of the results only trended positive and did not reach the threshold for strong evidence. This suggests the data, especially for the IIT predictions, may not be suited to draw strong conclusions.

### Data consistent with GNWT prediction of an onset and offset ignition

GNWT posits that a late, non-linear response (an ignition) amplifies and broadcasts information through long-range feedback from the prefrontal cortex to higher sensory areas. This ignition is proposed to occur within 300–500ms of both stimulus onset and offset. While the original Cogitate findings only observed evidence for the former, in our analysis, we observed findings consistent with both the onset and offset ignition.

The prediction of an ignition at stimulus offset represents a key point of divergence between GNWT and other theories of consciousness. This prediction may be considered a “bold” prediction of the theory (Negro, 2024); which are valuable because these predictions are unlikely to emerge in the data if the theory is false. While it seems counterintuitive that the brain would transmit signals from the PFC to higher-sensory regions after a stimulus is no longer present, GNWT predicts an ignition event after the offset of the current percept as it marks an update of the workspace to the new content (e.g., the fixation screen). Indeed, the presence of this response, while we cannot be certain that it is directly related to consciousness per se (see discussion below), is a unique prediction from GNWT which has robustly held up in these analyses for both the discovery and validation datasets. In effect, the presence of a response consistent with an offset ignition provides stronger support for the specific broadcasting mechanisms predicted by GWNT than the presence of an onset ignition alone.

It is worth addressing why a discrepancy has emerged between our findings and the previous analyses in relation to the presence of an offset ignition. One possibility is that the original Cogitate study focused on activity in the high-gamma band. This frequency-band was originally chosen because high-gamma band activity correlates with neural spiking (Manning et al., 2009). However, GNWT does accommodate for other spectral activity also. For instance, that distinct spectral signatures may underlie feedforward versus feedback signalling; such that bottom-up propagation is carried by gamma, and top-down broadcasting emerging primarily in the alpha-beta frequency range (Bastos et al., 2015; Mashour et al., 2020; Mejias et al., 2016). Therefore, it remains consistent with the theory for spectral ranges outside of gamma to underlie ignition events. While DCM for evoked responses are not able to tease apart the distinct influences of different spectral ranges (see DCM for the complex cross spectra instead; Friston et al., 2012), by restricting their analysis to high-gamma, it is possible the Cogitate Consortium filtered out lower-band top-down broadcasting. In contrast, our DCM analysis utilized the full spectrum of the evoked response, potentially capturing the low-frequency bands (theta, alpha, beta) that the theory identifies as important for feedback and global updating.

It is also important to note that, unlike the IIT analyses, the GNWT predictions tested here are less directly contingent on the network being uniquely face-selective. The finding that PFC feedback to higher-sensory regions is necessary to explain the data within the predicted 300–500ms windows remains broadly consistent with GNWT’s account, even if the observed connectivity patterns partly reflect general visual processing. Nonetheless, the absence of (strong) face specificity in the BMS validation analysis means we cannot be certain that the observed ignition uniquely reflects the broadcasting of face-specific conscious content, as opposed to a more generic updating of the workspace.

### Data consistent with IIT prediction of sustained FG connectivity

To evaluate the predictions of IIT, we first tested the necessity of sustained effective connectivity within a posterior cortical network. Specifically, testing whether feedback connections between the Fusiform Gyrus (FG) and V1 are necessary to explain the data throughout the stimulus duration. This is because IIT posits that the physical substrate of a conscious experience (the “main complex”) must exist as a maxima of irreducible integrated information, typically localized to a posterior “hot zone”. Unlike the phasic “ignition” of GNWT, this cause-effect structure must be sustained for the duration of the experience.

This first test challenged the hypothesis that the feedback connections from higher category selective areas to lower areas was needed to explain the data. In both the discovery and validation datasets, while these connections were necessary to explain the data in some time windows, this effect was not sustained across the entire stimulus duration.

Interestingly, this failure to find sustained connectivity between early visual areas and the FG replicates the findings of the original Cogitate analyses. The Consortium similarly observed that synchrony between early visual areas and category-selective areas was early and transient, challenging the theory’s prediction of sustained posterior synchrony. This failure to observe connectivity between these regions across both sets of analyses provides converging evidence that the absence of a sustained effect is likely not due to other factors (e.g. the high gamma-band frequencies analysed, as with GNWT) and suggests that these sustained responses are likely not present in the data.

Notably, in our tests we observed substantial volatility in the posterior probabilities (e.g., the sharp drops in evidence at time windows ending at 600ms and 700ms followed by a rebound at later windows in the discovery batch). We first sought to rule out fundamental inversion failures at the first-level DCMs. Our assessments confirmed that the mean free energy (an approximation of model evidence) behaved as expected for both discovery and validation batches, indicating a stable model convergence (that is, the dips in the group-level posterior probabilities did not correspond to dips in free energy at the first-level DCMs; see Figure 7).

We subsequently examined the temporal evolution of the group-level connectivity strengths against the main results for prediction one (Figure 7C and D). This inspection revealed that the group-level connectivity strengths (and their credibility intervals) shrank towards zero during time windows where the posterior probabilities dropped. Subsequently, an inspection of the connectivity parameters at some later time windows revealed that a late-stage rebound in posterior probability was primarily driven by a strong decrease in connectivity. It is beyond the scope of the present study to interpret the directions of these effects and why they occurred in this dataset, especially given that our continuous viewing paradigm necessarily entails a mixture of cognitive processing. However, the overarching interpretation is that the volatility of this parameter across the expanding time windows challenges the prediction that there is sustained, continuous connectivity between the FG and V1 in this data. Accordingly, the addition of small temporal increments can drastically alter the relative model evidence of the tested model.

For the second IIT prediction, we also evaluated the stability of intrinsic connectivity within the FG nodes. This analysis may be considered a more direct test of IIT, given its prediction of flexibility of the precise spatiotemporal boundaries of the ’main complex’, which is capable of expanding, contracting, or shifting based on the specific conscious contents (Tononi & Koch, 2015). Therefore, because it is possible the posterior network we tested above did not overlap entirely with the actual main complex predicted by IIT to underlie conscious perception, we focused on the FG, which is considered an established substrate of conscious face perception (Koch et al., 2016). We reasoned that the underlying main complex must necessarily invoke this substrate within its network, irrespective of the broader network it is a part of.

The discovery dataset showed mostly sustained connectivity throughout the stimulus duration. Although the posterior probability of the intrinsic parameter fluctuated and briefly dropped to an inconclusive level in one time window, it rebounded to strong evidence in subsequent windows, concluding with positive evidence by the final time window. The independent validation set confirmed this sustained response, exhibiting strong evidence of sustained intrinsic connectivity throughout the entire stimulus duration.

It is important to stress that the strength of this evidence is contingent upon the specificity of these findings to conscious face processing. This is because IIT predicts that the neural substrate of experience corresponds to the precise phenomenological content of the conscious percept — in this case, faces. Accordingly, a posterior network that responds most robustly to faces should explain face connectivity better than a network that responds most to objects. Whilst there was positive evidence that our BMS analysis favoured the face-specific network over the object network at most time windows, this preference did not reach the threshold for strong evidence at any time window across both datasets. Accordingly, while the broader pattern of sustained intrinsic connectivity within the FG is consistent with IIT, strong claims cannot be made from this data that these effects were unique to face-specific conscious processing.

The absence of face-selectivity in this data based on the BMS results may be related to two factors. First, MEG has relatively limited spatial resolution (compared to other techniques, such as fMRI). Though the MEG dataset was chosen here specifically for its excellent temporal resolution to test key spatiotemporal predictions, there may have been spatial leakage in the source estimates. The consequence is that resulting source estimates for both the face- and object-specific DCMs may be relatively similar even if the latent neural dynamics are face-selective. Second, while the FFA sits within the FG, the broader FG is not exclusively a face-selective region and has been shown to respond to a range of complex visual categories, including objects, bodies, and words, with selectivity emerging from fine-grained subregional organisation (Binder et al., 2009; Peelen & Downing, 2005; Weiner & Grill-Spector, 2012; Weiner & Zilles, 2016). The spatial proximity of these higher category-selective and face-selective subregions, in combination with the relatively lower spatial resolution of the MEG, may have contributed to the object-specific DCM network being difficult to strongly dissociate from the face-specific network.

The transient dip in evidence at the 600ms window in the discovery set could be related to the local maxima problem. Because DCM employs variational Laplace to iteratively optimize free energy (Friston et al., 2007), the algorithm can occasionally become trapped in a local maximum in this optimisation process, rather than the global maxima. However, given the subsequent recovery of these parameters, coupled with the replication in the validation cohort, it is plausible this dip reflects the algorithm converging on a non-optimal solution at this time window specifically, and not the others tested. This is a challenge inherent to our expanding time window approach, and we discuss this limitation in more detail below.

### Functional Connectivity vs Effective Connectivity

One intriguing aspect of these analyses are the fact that they diverge from some of the key findings from the Cogitate Consortium’s original analyses of the data. Specifically, while the original study presented a mixed verdict, finding partial support for both theories but failing to robustly observe the “ignition” or “sustained maintenance” predictions in full, our DCM analysis yielded broad, robust support for GNWT predictions and a more nuanced, yet still broadly consistent results with IIT.

This divergence may stem from the fundamental difference between functional and effective connectivity. The Cogitate Consortium extensively analysed the data using functional connectivity measures. At its core, these metrics quantify statistical dependencies rather than directed influences. Consequently, they are susceptible to confounds, where correlations between two regions can be driven by a third source rather than direct interaction, or by fluctuations in signal-to-noise ratios independent of coupling changes between the ROIs (Friston, 2011). In contrast, DCM assesses effective connectivity, the directed influence one neural system exerts over another. By explicitly modelling the underlying neuronal architecture and distinguishing feedback from feedforward connectivity, DCM is able to investigate the directed influences underlying broadcasting and maintenance that correlational measures may not be able to directly assess.

### Limitations

A primary limitation concerns the content-specificity of the network architecture. Although the DCM network was anchored to the FG on the basis of its established role in face perception (Kanwisher et al., 1997), and the face-specific network broadly had positive evidence over the object-specific network in the BMS analysis, this preference did not reach the threshold for strong evidence (PEP > 0.95) at any time window. Consequently, the possibility that the observed connectivity patterns, particularly the sustained intrinsic FG connectivity interpreted in relation to IIT, may reflect more general visual processing common to multiple stimulus categories cannot be definitively excluded. Future studies employing within-subject paradigms that directly contrast seen versus unseen stimuli, or that include null-stimulus baselines, would afford a more rigorous test of content-specificity.

A further methodological constraint of our temporal analyses relates to the GNWT prediction of a transient ignition approximately occurring within 300 to 500ms post-onset and offset. However, DCM for evoked responses evaluates network connectivity within an entire window and requires a fixed starting point at time zero to capture the initial sensory drive that perturbs the network (David et al., 2006). Consequently, our expanding window approach evaluated network dynamics across the broader 0 to 500ms epoch, rather than isolating the 300 to 500ms window as an independent window. Specifically, because the later neural dynamics are dependent on the initial timepoints, we cannot definitively claim that connections at the time windows ending at 300-500ms are uniquely necessary to explain the data; independent of the preceding time windows.

Similarly, this expanding time-window approach also adds caveats to the interpretation of our IIT tests. Specifically, because each successive time window is accumulative (i.e. includes data from preceding windows) and because DCM fits a generative model to the entire evoked response waveform, we cannot rule out that the resulting posterior estimates are strongly weighted by early, high-amplitude evoked responses at stimulus onset and remain inflated in later time windows primarily due to activity early time windows. Put differently, genuine sustained connectivity in late time windows may be indistinguishable from phasic connectivity that is driven by strong early evoked responses.

However, in both cases, if the models tested required a reduction in connectivity within the tested time windows to fit the data, this would indicate the model cannot support sustained processing or late-stage ignition under any parameterization. Therefore, while a negative result (i.e. a reduction in posterior probability at the tested time windows) would conclusively challenge either theory, a positive result (i.e. strong posterior probability at the tested time windows), as demonstrated here, should be interpreted with caution. Thus, our tests in principle can only convincingly refute the theories but not unequivocally support them.

An additional limitation concerns the robustness of the DCM inversion scheme, especially when inverted overlapping data, as is the case with our expanding time window approach. The inversion of DCM relies on the variational Laplace, an optimization regime that seeks to iteratively maximize the negative free energy (an approximation for the log model evidence). However, this optimization is non-trivial because the free energy landscape often contains multiple local maxima. Consequently, this carries a risk of converging on a local rather than global maximum (Daunizeau et al., 2011). Since our analysis involved inverting models across expanding time windows independently, this risk is amplified. That is, the optimization scheme may converge on different local solutions for adjacent time windows. Although, most of our results replicated across batches, it could be that instable estimates explains the few discrepant patterns of results; e.g. the slight differences between the discovery and validation datasets for the posterior (FG-V1) test. Specifically, the failure to replicate the connectivity pattern across these datasets may instead be the result of some time windows not converging upon the global maxima of free energy. In this scenario, the observed divergence would reflect a limitation of the optimization algorithm, rather than a genuine instability of the underlying effect.

A broader clarifying point is that the predictions tested from either theory were not mutually exclusive. Instead, we aimed to determine whether the predicted pattern of results for a given theory was present in the data. Consequently, we did not pit the theories against each other in a single model space. Such a comparison would be inappropriate for this dataset, primarily because the neural signal contains a mixture of conscious processing, sensory encoding, and other cognitive processing (such as attention, insofar as these processes are considered separable; see Koch & Tsuchiya, 2007; Pitts et al., 2018). Therefore, here we were unable to compare the relative model evidence of each theory.

## Conclusion

In conclusion, this study adjudicates between two leading theories of consciousness, IIT and GNWT, by investigating effective connectivity changes during conscious face perception. The results were consistent with both predictions of GNWT and one prediction of IIT, though these findings must be interpreted with important caveats due to a post-hoc validation analysis failing to show strong evidence that the networks tested were specific for faces. For our tests of GNWT, we identified robust evidence consistent with a late prefrontal ignition at stimulus onset and also at offset, the latter finding not observed by the Cogitate Consortium’s original analyses. For IIT, while the initial test of posterior feedback did not show sustained connectivity, challenging a key IIT prediction, we observed intrinsic connectivity consistent with a sustained response within the FG. However, as a follow-up BMS validation analysis did not reach the threshold for strong evidence, this finding should be interpreted cautiously as the data may not be well-suited to draw firm conclusions specific to face-processing. Additionally, as our time windows are cumulative, we cannot rule out that sustained intrinsic connectivity observed for IIT predictions or late prefrontal ignition observed for GNWT, are at least partially driven by strong connectivity in earlier time windows, which may drive neural dynamics at later (tested) time windows.

Ultimately, this study highlights the importance of falsification-based frameworks in the neuroscience of consciousness. By design, both the experimental design and our analytical approach shared an epistemic asymmetry, such that they were primarily designed to challenge theories rather than confirming them. Because positive evidence must be interpreted cautiously in this context (due to the absence of an unseen baseline condition and the cumulative nature of our expanding time windows) our models are (in principle) most informative when demonstrating that a predicted causal mechanism was statistically absent. While DCM is a rigorous tool for testing *a priori* hypotheses, its utility depends on the degree to which these hypotheses are precise. Continuing to translate broad theoretical constructs into testable, directed network models will be fruitful for elucidating the spatiotemporal mechanism underlying consciousness.

## Code availability

https://github.com/kbandara/cogitate_dcm

## Acknowledgements

The authors thank Karl Friston, Peter Zeidman, Pranay Yadav and the members of the SPM methods group, Lucia Melloni’s research group, and Heleen Slagter’s research group for their helpful discussions and feedback. We are also grateful to the Cogitate Consortium for collecting this data and making it publicly available. This research was supported by The University of Melbourne’s Research Computing Services and the Petascale Campus Initiative. This research was supported by the Commonwealth through an Australian Government Research Training Program Scholarship (DOI: https://doi.org/10.82133/C42F-K220) and the Melbourne School of Psychological Sciences PhD Studentship.

## References

1. Amunts, K., Malikovic, A., Mohlberg, H., Schormann, T., & Zilles, K. (2000). Brodmann’s areas 17 and 18 brought into stereotaxic space—where and how variable?. Neuroimage, 11(1), 66–84.

2. Amunts, K., Schleicher, A., Bürgel, U., Mohlberg, H., Uylings, H. B., & Zilles, K. (1999). Broca’s region revisited: cytoarchitecture and intersubject variability. Journal of Comparative Neurology, 412(2), 319–341.

3. Amunts, K., Weiss, P. H., Mohlberg, H., Pieperhoff, P., Eickhoff, S., Gurd, J. M., … & Zilles, K. (2004). Analysis of neural mechanisms underlying verbal fluency in cytoarchitectonically defined stereotaxic space—the roles of Brodmann areas 44 and 45. Neuroimage, 22(1), 42–56.

4. Ashburner, J., Barnes, G., Chen, C. C., Daunizeau, J., Flandin, G., Friston, K., … & Penny, W. (2014). SPM12 manual. *Wellcome Trust Centre for Neuroimaging, London*, UK, 2464(4), 53.

5. Bludau, S., Eickhoff, S. B., Mohlberg, H., Caspers, S., Laird, A. R., Fox, P. T., … & Amunts, K. (2014). Cytoarchitecture, probability maps and functions of the human frontal pole. Neuroimage, 93, 260–275.

6. Caspers, J., Zilles, K., Eickhoff, S. B., Schleicher, A., Mohlberg, H., & Amunts, K. (2013). Cytoarchitectonical analysis and probabilistic mapping of two extrastriate areas of the human posterior fusiform gyrus. Brain Structure and Function, 218(2), 511–526.

7. Eickhoff, S. B., Stephan, K. E., Mohlberg, H., Grefkes, C., Fink, G. R., Amunts, K., & Zilles, K. (2005). A new SPM toolbox for combining probabilistic cytoarchitectonic maps and functional imaging data. Neuroimage, 25(4), 1325–1335.

8. Ferrante, O., Liu, L., Minarik, T., Gorska, U., Ghafari, T., Luo, H., & Jensen, O. (2022). FLUX: A pipeline for MEG analysis. NeuroImage, 253, 119047.

9. Geyer, S. (2012). The microstructural border between the motor and the cognitive domain in the human cerebral cortex.

10. Geyer, S., Ledberg, A., Schleicher, A., Kinomura, S., Schormann, T., Bürgel, U., … & Roland, P. E. (1996). Two different areas within the primary motor cortex of man. Nature, 382(6594), 805–807.

11. Gramfort, A., Luessi, M., Larson, E., Engemann, D. A., Strohmeier, D., Brodbeck, C., … & Hämäläinen, M. S. (2014). MNE software for processing MEG and EEG data. NeuroImage, 86, 446–460.

12. Henssen, A., Zilles, K., Palomero-Gallagher, N., Schleicher, A., Mohlberg, H., Gerboga, F., … & Amunts, K. (2016). Cytoarchitecture and probability maps of the human medial orbitofrontal cortex. Cortex, 75, 87–112.

13. Lorenz, S., Weiner, K. S., Caspers, J., Mohlberg, H., Schleicher, A., Bludau, S., … & Amunts, K. (2017). Two new cytoarchitectonic areas on the human mid-fusiform gyrus. Cerebral Cortex, 27(1), 373–385.

14. Mattout, J., Henson, R. N., & Friston, K. J. (2007). Canonical source reconstruction for MEG. Computational Intelligence and Neuroscience, 2007(1), 067613.

15. Nolte, G. (2003). The magnetic lead field theorem in the quasi-static approximation and its use for magnetoencephalography forward calculation in realistic volume conductors. Physics in Medicine & Biology, 48(22), 3637.

16. Overgaard, M., & Sandberg, K. (2021). The Perceptual Awareness Scale—recent controversies and debates. Neuroscience of Consciousness, 2021(1), niab044.

17. Tononi, G. (2004). An information integration theory of consciousness. BMC Neuroscience, 5(1), 42.

18. Albantakis, L., Barbosa, L., Findlay, G., Grasso, M., Haun, A. M., Marshall, W., Mayner, W. G. P., Zaeemzadeh, A., Boly, M., Juel, B. E., Sasai, S., Fujii, K., David, I., Hendren, J., Lang, J. P., & Tononi, G. (2023). Integrated information theory (IIT) 4.0: Formulating the properties of phenomenal existence in physical terms. PLOS Computational Biology, 19(10), e1011465. 10.1371/journal.pcbi.1011465

19. Bastos, A. M., Vezoli, J., Bosman, C. A., Schoffelen, J.-M., Oostenveld, R., Dowdall, J. R., De Weerd, P., Kennedy, H., & Fries, P. (2015). Visual Areas Exert Feedforward and Feedback Influences through Distinct Frequency Channels. Neuron, 85(2), 390–401. 10.1016/j.neuron.2014.12.018

20. Binder, J. R., Desai, R. H., Graves, W. W., & Conant, L. L. (2009). Where Is the Semantic System? A Critical Review and Meta-Analysis of 120 Functional Neuroimaging Studies. Cerebral Cortex, 19(12), 2767–2796. 10.1093/cercor/bhp055

21. Daunizeau, J., David, O., & Stephan, K. E. (2011). Dynamic causal modelling: A critical review of the biophysical and statistical foundations. NeuroImage, 58(2), 312–322. 10.1016/j.neuroimage.2009.11.062

22. David, O., Kiebel, S. J., Harrison, L. M., Mattout, J., Kilner, J. M., & Friston, K. J. (2006). Dynamic causal modeling of evoked responses in EEG and MEG. NeuroImage, 30(4), 1255–1272. 10.1016/j.neuroimage.2005.10.045

23. Dehaene, S., & Changeux, J.-P. (2005). Ongoing Spontaneous Activity Controls Access to Consciousness: A Neuronal Model for Inattentional Blindness. PLOS Biology, 3(5), e141. 10.1371/journal.pbio.0030141

24. Dehaene, S., & Naccache, L. (2001). Towards a cognitive neuroscience of consciousness: Basic evidence and a workspace framework. *Cognition*, The Cognitive Neuroscience of Consciousness, 79(1), 1–37. 10.1016/S0010-0277(00)00123-2

25. Dehaene, S., Sergent, C., & Changeux, J.-P. (2003). A neuronal network model linking subjective reports and objective physiological data during conscious perception. Proceedings of the National Academy of Sciences, 100(14), 8520–8525. 10.1073/pnas.1332574100

26. Friston, K. J. (2011). Functional and Effective Connectivity: A Review. Brain Connectivity, 1(1), 13–36. 10.1089/brain.2011.0008

27. Friston, K. J., Bastos, A., Litvak, V., Stephan, K. E., Fries, P., & Moran, R. J. (2012). DCM for complex-valued data: Cross-spectra, coherence and phase-delays. *NeuroImage*, Neuroergonomics: The Human Brain in Action and at Work, 59(1), 439–455. 10.1016/j.neuroimage.2011.07.048

28. Friston, K. J., Harrison, L., & Penny, W. (2003). Dynamic causal modelling. NeuroImage, 19(4), 1273–1302. 10.1016/S1053-8119(03)00202-7

29. Garrido, M. I., Kilner, J. M., Kiebel, S. J., & Friston, K. J. (2007). Evoked brain responses are generated by feedback loops. Proceedings of the National Academy of Sciences, 104(52), 20961–20966. 10.1073/pnas.0706274105

30. Hyvarinen, A. (1999). Fast and robust fixed-point algorithms for independent component analysis. IEEE Transactions on Neural Networks, 10(3), 626–634. 10.1109/72.761722

31. Jansen, B. H., & Rit, V. G. (1995). Electroencephalogram and visual evoked potential generation in a mathematical model of coupled cortical columns. Biological Cybernetics, 73(4), 357–366. 10.1007/BF00199471

32. Kanwisher, N., McDermott, J., & Chun, M. M. (1997). The fusiform face area: A module in human extrastriate cortex specialized for face perception. The Journal of Neuroscience: The Official Journal of the Society for Neuroscience, 17(11), 4302– 4311. 10.1523/JNEUROSCI.17-11-04302.1997

33. Kass, R. E., & Raftery, A. E. (1995). Bayes Factors. Journal of the American Statistical Association, 90(430), 773–795. 10.1080/01621459.1995.10476572

34. Kiebel, S. J., Garrido, M. I., & Friston, K. J. (2007). Dynamic causal modelling of evoked responses: The role of intrinsic connections. NeuroImage, 36(2), 332–345. 10.1016/j.neuroimage.2007.02.046

35. Koch, C., Massimini, M., Boly, M., & Tononi, G. (2016). Neural correlates of consciousness: Progress and problems. Nature Reviews Neuroscience, 17(5), Article 5. 10.1038/nrn.2016.22

36. Koch, C., & Tsuchiya, N. (2007). Attention and consciousness: Two distinct brain processes. Trends in Cognitive Sciences, 11(1), 16–22. 10.1016/j.tics.2006.10.012

37. Litvak, V., Mattout, J., Kiebel, S., Phillips, C., Henson, R., Kilner, J., Barnes, G., Oostenveld, R., Daunizeau, J., Flandin, G., Penny, W., & Friston, K. (2011). EEG and MEG Data Analysis in SPM8. Computational Intelligence and Neuroscience, 2011(1), 852961. 10.1155/2011/852961

38. Manning, J. R., Jacobs, J., Fried, I., & Kahana, M. J. (2009). Broadband shifts in local field potential power spectra are correlated with single-neuron spiking in humans. The Journal of Neuroscience: The Official Journal of the Society for Neuroscience, 29(43), 13613–13620. 10.1523/JNEUROSCI.2041-09.2009

39. Mashour, G. A., Roelfsema, P., Changeux, J.-P., & Dehaene, S. (2020). Conscious Processing and the Global Neuronal Workspace Hypothesis. Neuron, 105(5), 776–798. 10.1016/j.neuron.2020.01.026

40. Mejias, J. F., Murray, J. D., Kennedy, H., & Wang, X.-J. (2016). Feedforward and feedback frequency-dependent interactions in a large-scale laminar network of the primate cortex. Science Advances, 2(11), e1601335. 10.1126/sciadv.1601335

41. Melloni, L. (2022). On keeping our adversaries close, preventing collateral damage, and changing our minds. Comment on Clark et al. Journal of Applied Research in Memory and Cognition, 11(1), 45–49. 10.1037/mac0000009

42. Melloni, L., Mudrik, L., Pitts, M., Bendtz, K., Ferrante, O., Gorska, U., Hirschhorn, R., Khalaf, A., Kozma, C., Lepauvre, A., Liu, L., Mazumder, D., Richter, D., Zhou, H., Blumenfeld, H., Boly, M., Chalmers, D. J., Devore, S., Fallon, F., … Tononi, G. (2023). An adversarial collaboration protocol for testing contrasting predictions of global neuronal workspace and integrated information theory. PLOS ONE, 18(2), e0268577. 10.1371/journal.pone.0268577

43. Negro, N. (2024). (Dis)confirming theories of consciousness and their predictions: Towards a Lakatosian consciousness science. Neuroscience of Consciousness, 2024(1), niae012. 10.1093/nc/niae012

44. Peelen, M. V., & Downing, P. E. (2005). Selectivity for the Human Body in the Fusiform Gyrus. Journal of Neurophysiology, 93(1), 603–608. 10.1152/jn.00513.2004

45. Pitts, M. A., Lutsyshyna, L. A., & Hillyard, S. A. (2018). The relationship between attention and consciousness: An expanded taxonomy and implications for ‘no-report’ paradigms. Philosophical Transactions of the Royal Society B: Biological Sciences, 373(1755), 20170348. 10.1098/rstb.2017.0348

46. Poldrack, R. A., Baker, C. I., Durnez, J., Gorgolewski, K. J., Matthews, P. M., Munafò, M. R., Nichols, T. E., Poline, J.-B., Vul, E., & Yarkoni, T. (2017). Scanning the horizon: Towards transparent and reproducible neuroimaging research. Nature Reviews Neuroscience, 18(2), 115–126. 10.1038/nrn.2016.167

47. Stephan, K. E., Penny, W. D., Daunizeau, J., Moran, R. J., & Friston, K. J. (2009). Bayesian model selection for group studies. NeuroImage, 46(4), 1004–1017. 10.1016/j.neuroimage.2009.03.025

48. Stephan, K. E., Penny, W. D., Moran, R. J., den Ouden, H. E. M., Daunizeau, J., & Friston, K. J. (2010). Ten simple rules for dynamic causal modeling. Neuroimage, 49(4), 3099– 3109. 10.1016/j.neuroimage.2009.11.015

49. Taulu, S., & Kajola, M. (2005). Presentation of electromagnetic multichannel data: The signal space separation method. Journal of Applied Physics, 97, 124905–124905–124910. 10.1063/1.1935742

50. Tononi, G., Boly, M., Massimini, M., & Koch, C. (2016). Integrated information theory: From consciousness to its physical substrate. Nature Reviews Neuroscience, 17(7), Article 7. 10.1038/nrn.2016.44

51. Tononi, G., & Koch, C. (2015). Consciousness: Here, there and everywhere? Philosophical Transactions of the Royal Society B: Biological Sciences, 370(1668), 20140167. 10.1098/rstb.2014.0167

52. Weiner, K. S., & Grill-Spector, K. (2012). The improbable simplicity of the fusiform face area. Trends in Cognitive Sciences, 16(5), 251–254. 10.1016/j.tics.2012.03.003

53. Weiner, K. S., & Zilles, K. (2016). The anatomical and functional specialization of the fusiform gyrus. Neuropsychologia, 83, 48–62. 10.1016/j.neuropsychologia.2015.06.033

